# SMRT-AgRenSeq-d in potato (Solanum tuberosum) identifies candidates for the nematode resistance Gpa5

**DOI:** 10.1101/2022.12.08.519582

**Authors:** Yuhan Wang, Lynn H Brown, Thomas M Adams, Yuk Woon Cheung, Jie Li, Vanessa Young, Drummond T Todd, Miles R Armstrong, Konrad Neugebauer, Amanpreet Kaur, Brian Harrower, Stan Oome, Xiaodan Wang, Micha Bayer, Ingo Hein

## Abstract

Potato is the third most important food crop in the world. Diverse pathogens threaten sustainable crop production but can be controlled, in many cases, through the deployment of disease resistance genes belonging to the family of nucleotide-binding, leucine-rich-repeat (NLR) genes.

To identify functional NLRs in established varieties, we have successfully established SMRT-AgRenSeq in tetraploid potatoes and have further enhanced the methodology by including dRenSeq in an approach that we term SMRT-AgRenSeq-d. The inclusion of dRenSeq enables the filtering of candidates after the association analysis by establishing a presence/absence matrix across resistant and susceptible potatoes that is translated into an F1 score. Using a SMRT-RenSeq based sequence representation of the NLRome from the cultivar Innovator, SMRT-AgRenSeq-d analyses reliably identified the late blight resistance benchmark genes *R1, R2-like, R3a* and *R3b* in a panel of 117 varieties with variable phenotype penetrations. All benchmark genes were identified with an F1 score of 1 which indicates absolute linkage in the panel.

When applied to the elusive nematode disease resistance gene *Gpa5* that controls the Potato Cyst Nematode (PCN) species *Globodera pallida* (pathotypes Pa2/3), SMRT-AgRenSeq-d identified nine strong candidates. These map to the previously established position on potato chromosome 5 and are potential homologs of the late blight resistance gene *R1*.

Assuming that NLRs are involved in controlling many types of resistances, SMRT-AgRenSeq-d can readily be applied to diverse crops and pathogen systems. In potato, SMRT-AgRenSeq-d lends itself, for example, to further study the elusive PCN resistances *H1* or *H3* for which phenotypic data exist.

## Introduction

Potato is the third most important global food crop and is both nutritionally and economically valuable [1]. Potato is a staple of many diets around the world, and diseases can thus have a significant impact on crop production and food security. Diverse and phylogenetically unrelated pathogens ranging from oomycetes, fungi, bacteria, viruses, nematodes, and insects can cause significant crop losses in potato [2].

The Potato Cyst Nematode (PCN) species *Globodera rostochiensis* and *G. pallida* are economically important pathogens of potato and are present in most potato growing regions of the world [3]. PCN is difficult to eradicate once established as cysts can survive for over 20 years in the soil, rendering even longer crop rotations insufficient for clearing land [4,5]. Thus, the impact of PCN goes beyond the immediate yield losses, which are estimated to be around 9% on average in susceptible potatoes, with the presence of PCN rendering land unsuitable for high-health potato seed production, which significantly impacts on the crop supply chain [4,6].

The only cloned disease resistance genes effective against PCN are *Gpa2* and *Gro1-4* which encode for canonical nucleotide-binding, leucine-rich-repeat (NLR) genes [7,8]. In current potato cultivars, the most effective resistance gene used to control *G. pallida* is *Gpa5*, which has been introduced from the wild Solanaceae species *Solanum vernei* [9]. The resistance has been mapped to potato chromosome 5 [9,10] and a haplotype specific PCR marker, HC, was developed which corresponds in its position to an NLR-rich locus on chromosome 5 [11]. This locus is also associated with the late blight resistance gene *R1,* amongst other functional NLRs such as *Grp1* and *H2* [3,12]. Intriguingly, Rouppe van der Voort *et al.* [9] and an independent study aimed at mapping an *S. vernei* source of resistance against *G. pallida*, independently revealed a further QTL, *Gpa6*, on chromosome 9 that complements the major effect mapped to chromosome 5 [13,14].

To specifically study potato NLRs, RenSeq was developed as a target enrichment tool following the annotation of NLRs in the reference genome DM1-3 516 R44 (DM) [15–17]. In combination with PacBio-based sequencing of long genomic DNA fragments, described as SMRT-RenSeq, the technology enables the representation of entire plant NLRomes [18,19]. A diagnostic version of RenSeq, dRenSeq, has been used to ascertain the presence/absence of known NLRs in wild species or cultivars. Thus, dRenSeq aids the identification of novel resistances in germplasm collections as well as the deployment and pyramiding of complementary resistances in cultivars [20,21].

RenSeq has also been adapted for association genetics (AgRenSeq) which takes advantage of independent recombination events that occur during meiosis. AgRenSeq, which determines the potential of candidate NLRs being responsible for resistance traits, uses *k*-mer based association and has initially been implemented in *Aegilops tauschii,* a wild progenitor of the D subgenome of hexaploid bread wheat [22]. AgRenSeq has also been performed with PacBio HiFi reads to identify candidate resistance genes in Rye controlling *Puccinia recondita* f. sp. *secalis* [23]. This approach was termed SMRT-AgRenSeq.

Here we apply AgRenSeq for the first time in potatoes to identify candidate NLRs that are associated with the *Gpa5* resistance linked to the HC marker. To study *Gpa5*, we selected the tetraploid potato cultivar Innovator as a reference, since this cultivar is known to exhibit high levels of *G. pallida* resistance based on the presence of *Gpa5* [3]. As previously shown, Innovator also contains four known functional NLRs effective against *P. infestans*: *R1*, *R2-like*, *R3a,* and a variant of *R3b* [20]. These four NLRs reside in three clusters of different sizes and thus function as ideal benchmark genes to test the accuracy of the NLR assembly and subsequent association studies.

We further developed SMRT-AgRenSeq by including a dRenSeq analysis to validate the association of candidates and, importantly, identify candidate NLRs that are 100% linked to the resistance phenotype. We refer to this combination as SMRT-AgRenSeq-d and suggest that this technology is well suited to identify resistances against diverse pathogens that can be controlled through functional NLRs [24].

## Results and Discussion

SMRT-AgRenSeq-d encompasses a representation of the NLRs from a plant with the desired resistance, a panel of plants with contrasting phenotypes to ascertain the *k*-mer based association, and dRenSeq-based assessment of candidates. The benchmark genes *Rpi-R1, Rpi-R2-like*, *Rpi-R3a* and *Rpi-R3b* in Innovator have been utilised to provide a quality check throughout the analysis.

### *De-novo* assembly of Illumina-based RenSeq reads fails to represent benchmark NLRs

To assess the accuracy of the NLR representation from Illumina sequences, we used RenSeq reads generated from the potato cultivar Innovator [20]. The *de novo* assembly of Illumina-based RenSeq reads yielded 9,756 contigs with an average length of 1,060.4 bp of which 1,111 contigs contained signatures of NLRs. NLR-Annotator [25] predicted 1,150 NLRs in this assembly (Supplementary Table 1). The integrity of the four benchmark genes was assessed in the assembled contigs (Table 1). Only *Rpi-R1* was accurately and completely represented in the contig scf7180000044808. In contrast, *Rpi-R2-like*, and the *Rpi-R3b* variant were partially and incorrectly represented in contigs scf7180000045065 and scf7180000045838, respectively. For *Rpi-R2-like*, more than 7% sequence diversity was observed compared to the reference, and for *Rpi-R3b,* 38% of the gene was missing. Representation of *Rpi-R3a* was split over two contigs, scf7180000044942 and scf7180000045673.

**Table 1.**
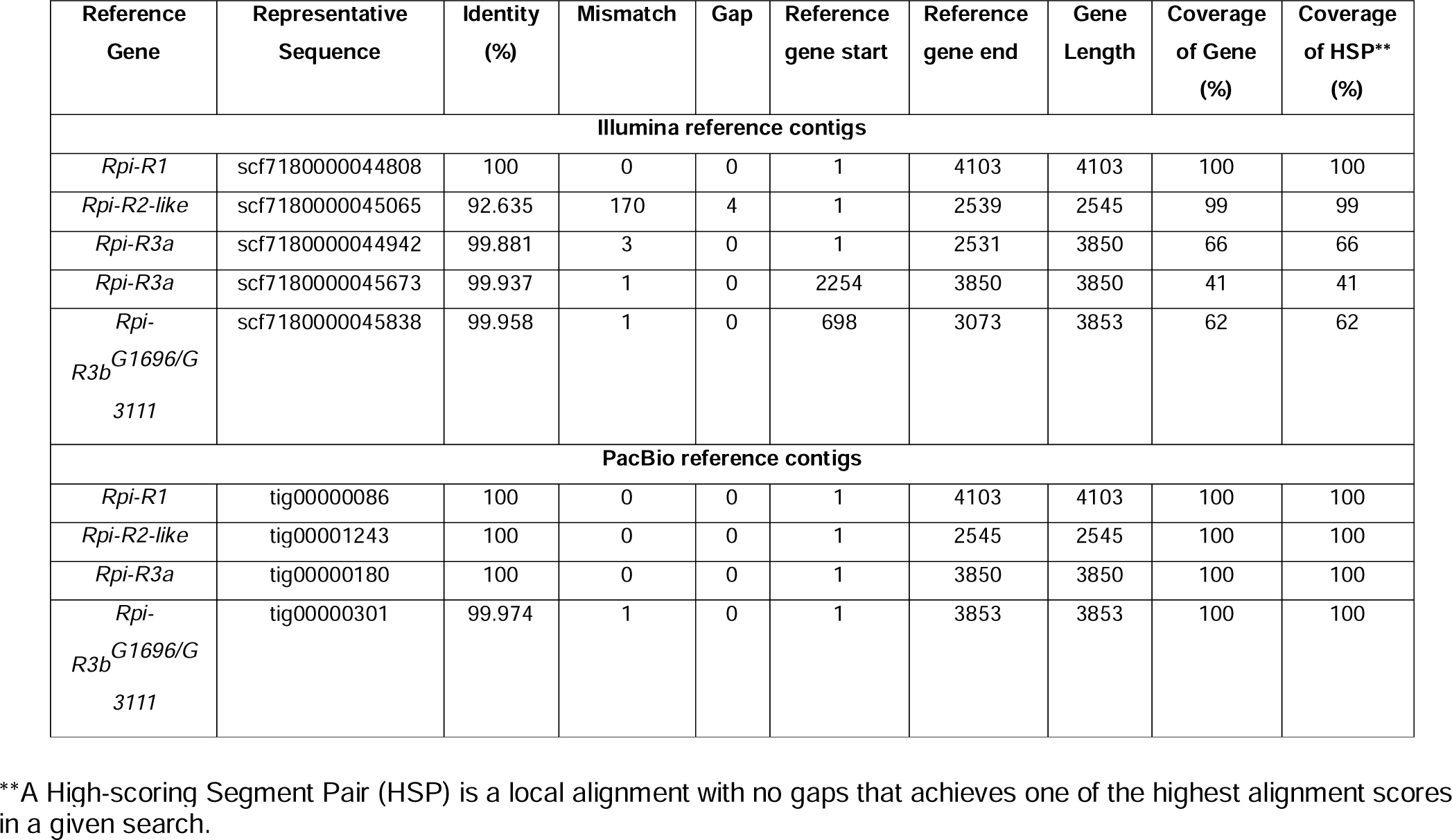
Representation of four benchmark genes in the potato cultivar Innovator based on Illumina and PacBio RenSeq read assemblies.

Considering that the average full-length NLR in potato is approximately 3 kb in length [16], the partial representation of *Rpi-R3a* and *Rpi-R3b* is consistent with the relatively small average contig size observed. Importantly, the 150 bp Illumina reads not only affect the assembly length of NLRs, but also their accuracy, the lack of which is pronounced in *Rpi-R2-like* and *Rpi-R3a/R3b.* These genes originate in the two largest NLR clusters in the potato genome [16]. The processes leading to larger homogeneous clusters containing highly similar NLRs are typically governed by tandem duplications [26]. It is likely that the assembly of the short reads collapsed NLR gene clusters and insufficiently distinguished between alleles and paralogs. Evidence for this was provided by comparing the number of predicted NLRs in the doubled monoploid potato reference clone DM and Innovator. DM contains 755 NLRs [15] whereas the Illumina-based assembly of NLRs from the tetraploid cultivar Innovator predicted 1,150, which suggests only a modest increase of 53% of the total number of NLRs.

### PacBio HiFi-based assembly of RenSeq reads generates a high-quality NLRome representation

The combination of RenSeq with single-molecule real-time (SMRT) sequencing (SMRT-RenSeq) has overcome the limitations of Illumina-based assemblies in the cloning of NLRs in wild diploid potato species [19]. The assembly of HiFi RenSeq reads from Innovator yielded 4,885 contigs with an average length of 11,279 bp. Amongst those, 1,760 contigs contained a total of 2,418 predicted NLRs (Supplementary Table 1).

In this assembly, the benchmark genes *Rpi-R1, Rpi-R2-like* and *Rpi-R3a* were represented completely and accurately in a single contig each (Table 1).

A comparison between the previously identified variant of *Rpi-R3b* in Innovator*, Rpi-R3b^G1696/G3111^*, with the well supported PacBio contig tig00000301 revealed 100% coverage of this gene and 99.974% identity to *Rpi-R3b^G1696/G3111^*with only one mismatch at position C918. We referred to this variant as *Rpi-R3b^C918/G1696/G3111^.* Interestingly, Illumina RenSeq reads from Innovator and other cultivars, except for Brodie, could not distinguish *Rpi-R3b^G1696/G3111^* from *Rpi-R3b^C918/G1696/G3111^* (Supplementary Table 2). This suggests that highly similar sequences obscure the additional sequence polymorphism, or that some cultivars contain both variants.

These results independently confirm and quantify the advances that SMRT-RenSeq offers for the highly accurate representation of the complex NLR gene family that cannot be achieved through Illumina-based assemblies [18,23,27]. In addition, these contigs, which are larger in size compared to the described Illumina-derived contigs, can contain multiple NLRs and their flanking regions on single contigs which aids functional analyses such as expression of candidates with their native regulatory elements.

A phylogenetic tree was constructed from the SMRT-RenSeq derived NLRs using the NB domains of predicted NLRs from Innovator. Further included in the tree is a selection of reference NLRs and two outgroup genes with NB domains from Humans (*Apaf-1*) and *C*. *elegans (Ced4)* (Figure 1; Supplementary Figure 1). The resulting tree shows clear groupings of NLRs into numerous distinct clades and emphasises the scale of duplications that have occurred in tetraploid potatoes. In addition to highlighting the NLRs related to *Rpi-R1, Rpi-R2-like, Rpi-R3a* and *Rpi-R3b*, a clear NRC (NLR Required for Cell death) clade [28] is apparent (Figure 1; Supplementary Figure 1).

**Figure 1:**
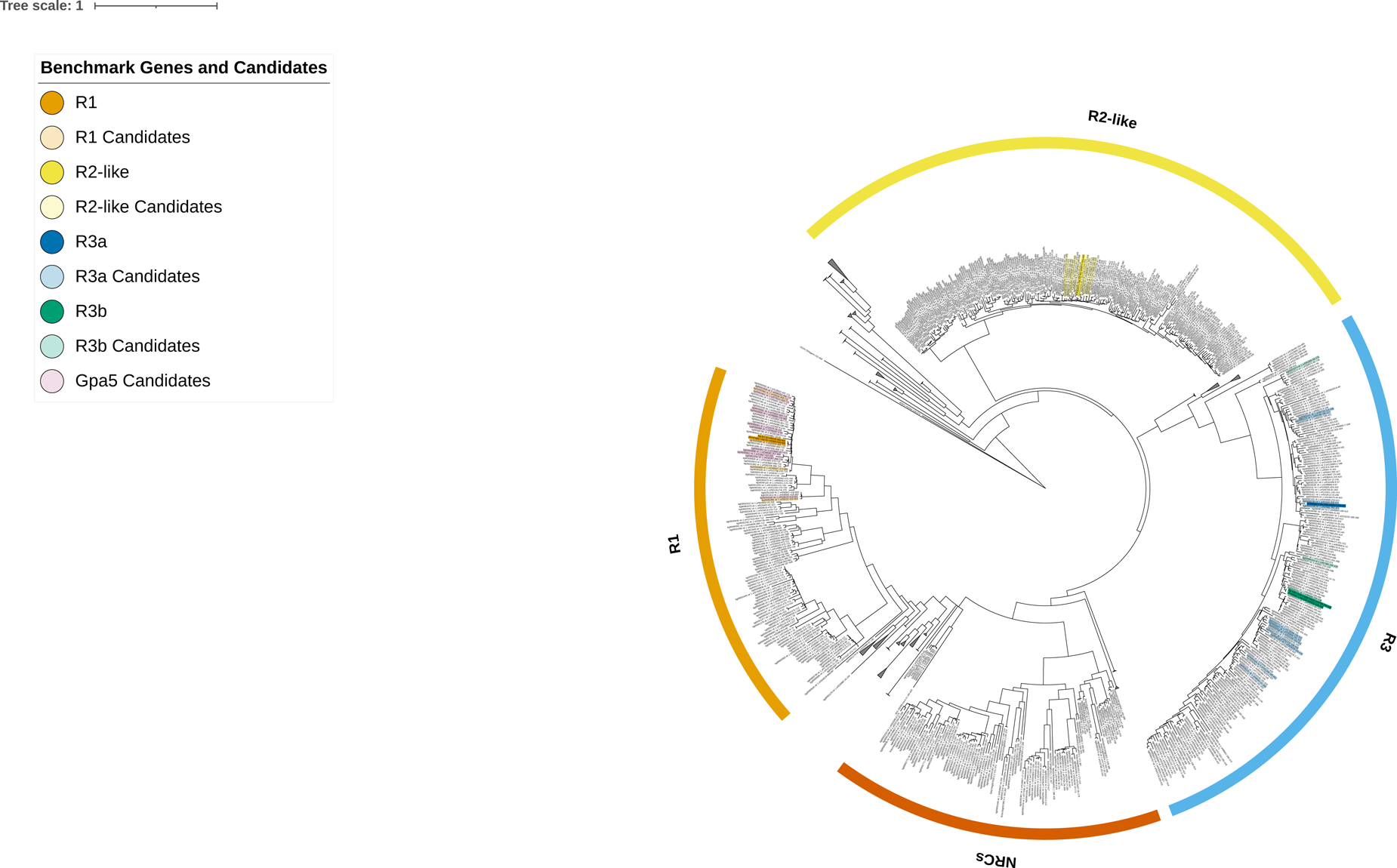
Phylogenetic tree displaying NB domains of Innovator NLRs following assembly of SMRT-RenSeq reads. Expanded are the clades containing *NRCs, Rpi-R1*, *Rpi-R2-like, Rpi-R3a* and *Rpi-R3b*. Collapsed clades either consist of automatic predictions from NLR-Annotator or of NLR clades not investigated in this study. Highlighted are the identified benchmark genes and candidates from each association with F1=1. The NB containing proteins APAF1 from *H*. *sapiens* and CED4 from *C*. *elegans* act as outgroups. The tree scale indicates substitutions/site.

### Development of an association panel

The cultivar Innovator was released in 1999 and has since been extensively used in breeding programs and is attributed as a direct parent to over 25 named varieties [29] (Supplementary Table 3). To study *Gpa5* and to take advantage of the wide genetic reach of Innovator, we developed an association panel of 117 potato accessions with contrasting resistance/susceptible phenotypes against *G. pallida*, associated with the presence/absence of *Gpa5.* Resistance scores were either determined by replicated canister tests or, if readily available publicly, extracted from databases (Supplementary Table 2). For *Gpa5*, the panel contained 23 highly resistant clones with a *G. pallida* resistance score of between 7 and 9 all of which tested positive for the presence of the HC marker, and 94 clones with a score of 1–3 (Supplementary Table 2).

Further, we assessed the same panel for the sequence representation of the late blight resistance genes *Rpi-R1, Rpi-R2-like, Rpi-R3a* and *Rpi-R3b* through dRenSeq, as they have been identified in Innovator previously [20]. The sequence representation took into consideration known family members with sequence polymorphisms. For example, in Armstrong *et al*. [20] we showed that Pentland Dell contains the *Rpi-R2* family member *Rpi-abpt^T86^* which has also been identified in the cultivar Bionica and the breeding clone 2573(2). Informed by the natural diversity of *Rpi-R2-like,* which in the cultivar Pentland Dell has 97.4% dRenSeq coverage [20], we set a minimum cut-off value of more than 97% sequence representation for a gene to be classified as ‘present’. This was converted into a presence/absence matrix for the four genes, which form the benchmarks for this study, across the association panel (Supplementary Table 2).

The representation of the benchmark genes in the panel was 10 clones for *Rpi-R2-like*, 34 for *Rpi-R1*, 44 for *Rpi-R3a* and 60 for *Rpi-R3b*. Interestingly, all clones that contained *Rpi-R3a* also contained *Rpi-R3b,* but not *vice versa*.

### SMRT-AgRenSeq-d development for *R3a* as a proof of concept

Since potato is an autotetraploid crop with complex tetrasomic inheritance patterns, we assessed first the suitability of SMRT-AgRenSeq-d on the benchmark genes *Rpi-R1, Rpi-R2-like, Rpi-R3a* and *Rpi-R3b* (Figure 2; Supplementary Figures 2 – 4).

**Figure 2:**
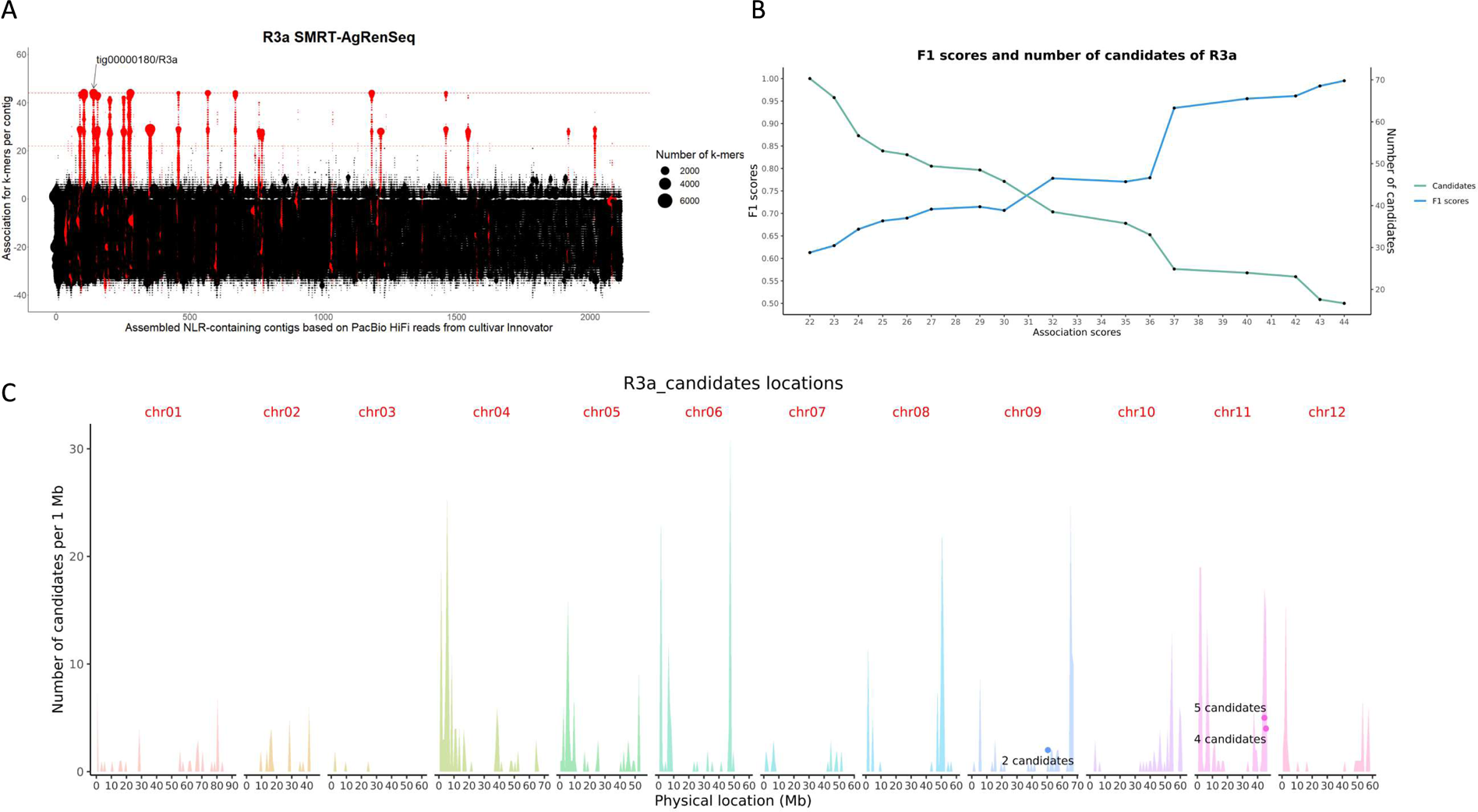
SMRT-AgRenSeq-d of *Rpi-R3a*. a) Identification of NLR containing contigs in the reference cultivar Innovator with association to *Rpi-R3a*. Columns on the X-axis represents NLR containing contigs, with dots indicating mapped *k*-mers and their association score to *Rpi-R3a* on the Y-axis. Dashed lines highlight highest association score (44, red) and chosen threshold (22, Orange). b) F1 scores and number of candidate NLRs for each association score group. c) The predicted positions of *Rpi-R3a* candidates with F1=1 are shown (dots) in relation to all NLRs from the reference DM v6.1 mapped over 12 chromosomes (peaks).

The SMRT-AgRenSeq-d study for *Rpi-R3a* yielded a maximum positive association score of 44 (Figure 2a). To systematically assess the resulting candidates and to be as inclusive as possible in the analysis, we selected half of the maximum association score as an initial cut-off for candidate selection. At this lower association score of 22, 46 contigs were identified that contain 73 candidate NLRs. The sequences of these 73 predicted NLRs were analysed by dRenSeq to independently validate their prevalence in the 44 clones that contain the reference sequence of *Rpi-R3a* and their absence in the 73 potato clones identified as missing *Rpi-R3a* (Supplementary Table 4). Based on this graphical genotyping depicting the presence/absence of candidate genes in the respective pools, an F1 score [30] was determined. An F1 score of 1 indicates the presence of the candidate gene in all clones from the positive panel and the consistent absence in all negative clones. Importantly, the *Rpi-R3a* containing contig tig00000180 (Table 1) was amongst these highly positively associated candidates (Supplementary Table 4).

A score of less than 1 therefore indicates false positives (e.g., candidates that are not present in all clones from the positive set) and/or false negatives (e.g., candidates that are present in the negative set). This conceptually equates to identifying recombination events in potato clones that have, for example, lost a specific candidate. Thus, the F1 score, which is determined through the dRenSeq analysis, complements the association score by confirming and further prioritising candidates. Interestingly, candidates that yielded an F1 score of less than 0.7, some of which had an association score of 37, displayed unsystematic presence/absence patterns that are inconsistent with robust association.

The analysis of candidates with an F1 score of larger than 0.7 but smaller than 1 highlighted examples of potential recombination and false positives, including two NLRs with an association score of 44 from contig tig00000196 in the cultivar Cammeo that does not contain *Rpi-R3a* (Supplementary Table 4). Consequently, their F1 scores were lower than 1 at 0.9887. *Rpi-R3b* was also present in the *Rpi-R3a* candidate gene list, on contig tig00000301 (Table 1, Supplementary Table 4). This is consistent with the finding that all *Rpi-R3a* containing clones also contained *Rpi-R3b* (Supplementary Table 2). However, as other potato clones also contain *Rpi-R3b,* yet do not contain *Rpi-R3a*, the F1 score for this contig was again less than 1. Similarly, five candidates from contigs tig00000247, tig00000325 and tig00000301 were missing in the cultivar Brodie that contains *Rpi-R3a* (Supplementary Table 4) and hence also had F1 scores below 1. Consistent with this analysis, we found that the number of candidates is inversely related to the F1 score (Figure 2b).

Only 11 putative NLRs had an F1 score of 1 and, as mentioned above, include contig tig00000180 that contains the full-length *Rpi-R3a* gene sequence alongside a closely related gene. We surmise that these 11 genes are physically linked and likely represent paralogous sequences, as alleles would have been genetically randomly distributed owing to the tetrasomic inheritance patterns of potato. Further, when mapped to the DM reference genome, nine out of the 11 candidates resided in the predicted *R3* locus on chromosome 11 (Figure 2c). The remaining two genes had their best BLASTn hit on DM chromosome 9 which contains sequences that belong to the *R3* family as demonstrated by Jupe *et al.* [16]. Consistent with this, all 11 candidates reside in the *Rpi-R3a/R3b* clade of the phylogenetic tree (Figure 1).

### SMRT-AgRenSeq-d validation for benchmark genes *Rpi-R3b, Rpi-R2-like and Rpi-R1*

Using the same SMRT-AgRenSeq-d approach for the benchmark genes *Rpi-R3b, Rpi-R2-like* and *Rpi-R1* independently validated our methodology and successfully identified the functional genes amongst the positively associated contigs and specifically amongst those yielding an F1 score of 1. Thus, the integration of dRenSeq to validate the association of NLRs and to determine the F1 score, is a valuable step in reducing the number of candidates rapidly.

Following the SMRT-AgRenSeq-d analyses for *Rpi-R3b,* 83 NLR containing contigs with 114 NLRs were identified with a minimum association score of 30 (Supplementary Figure 2). When filtered for an F1 score of larger than 0.7, 21 contigs that contained 31 NLRs were identified with an association score of between 56 and 60 (Supplementary Table 5). Owing to the absence of candidates in potatoes that contain *Rpi-R3b* and the presence of candidates in potatoes that do not contain *Rpi-R3b*, only three contigs remained with an F1 score of 1. These include tig00000194 and tig00000301 of which the latter contains *Rpi-R3b* (Table 1). Further, all genes map to the *R3* locus on potato chromosome 11 (Supplementary Figure 2c) and reside in the phylogenetic clade associated with *Rpi-R3b* (Figure 1). The *Rpi-R3a* containing contig tig00000180 yielded an association score of 59 but, since this gene was absent in 16 of the 60 potatoes with *Rpi-R3b*, the F1 score was reduced to approximately 0.85 which ruled out this contig as a candidate (Supplementary Table 5).

Representation of the *Rpi-R2* gene family in the association set was the lowest of all benchmark genes. Despite less than 10% penetration of the phenotype in the panel (10 out of 117; Supplementary Table 2), SMRT-AgRenSeq-d identified ten candidate NLRs with an F1 score of 1 which include the *Rpi-R2-like* containing contig tig00001243 (Supplementary Figure 3; Supplementary Table 6, Table 1). These candidates all map to the *Rpi-R2* locus on potato chromosome 4 (Supplementary Figure 3c) and are members of the *Rpi-R2* clade in the phylogenetic tree (Figure 1).

The validation of SMRT-AgRenSeq-d on the *Rpi-R1* benchmark gene yielded the lowest number of associated contigs (Supplementary Figure 4) which is most likely a reflection of the smaller size of the *Rpi-R1* gene cluster compared to *Rpi-R2* and *Rpi-R3a/b*. Indeed, in DM, *Rpi-R1* is one of the smallest clusters of NLRs [16]. At a minimum association score of 17, 16 NLR containing contigs containing 20 NLRs were identified (Supplementary Figure 4). After filtering for an F1 score of larger than 0.7, 11 contigs had association scores between 22 and 34 and those encode for 13 NLRs (Supplementary Table 7). The *Rpi-R1* containing contig, tig00000086 (Table 1) was successfully identified with an F1 score of 1.

### SMRT-AgRenSeq-d identified nine candidates for Gpa5

Following the development and independent validation of SMRT-AgRenSeq-d on all four benchmark genes, the approach was applied to identify candidates for the resistance gene *Gpa5*. The association panel of 117 potatoes, as described above, includes 23 highly resistant clones with a *G. pallida* resistance score of between 7 and 9 and which tested positive for the HC marker [11], alongside 94 susceptible clones with a resistance score of 1 – 3 (Supplementary Table 2).

Conducting SMRT-AgRenSeq-d on this panel of highly resistant and highly susceptible potatoes identified 78 NLR-containing contigs with 139 predicted NLRs and with a minimum association score of 12 (Figure 3a). Following dRenSeq analysis and filtering of candidates that exceed an F1 score of 0.7, 46 contigs encoding for 79 predicted NLRs were identified with an association score between 13 and 23 (Supplementary Table 8). These NLRs mapped to three positions in the potato genome, chromosome 5, chromosome 7, and chromosome 9 (Supplementary Table 8). The position on chromosome 5 is consistent with previous mapping studies conducted by Rouppe van der Voort *et al.* [9] which also highlighted the position on chromosome 9 that was also observed by Bryan *et al.* [13].

**Figure 3:**
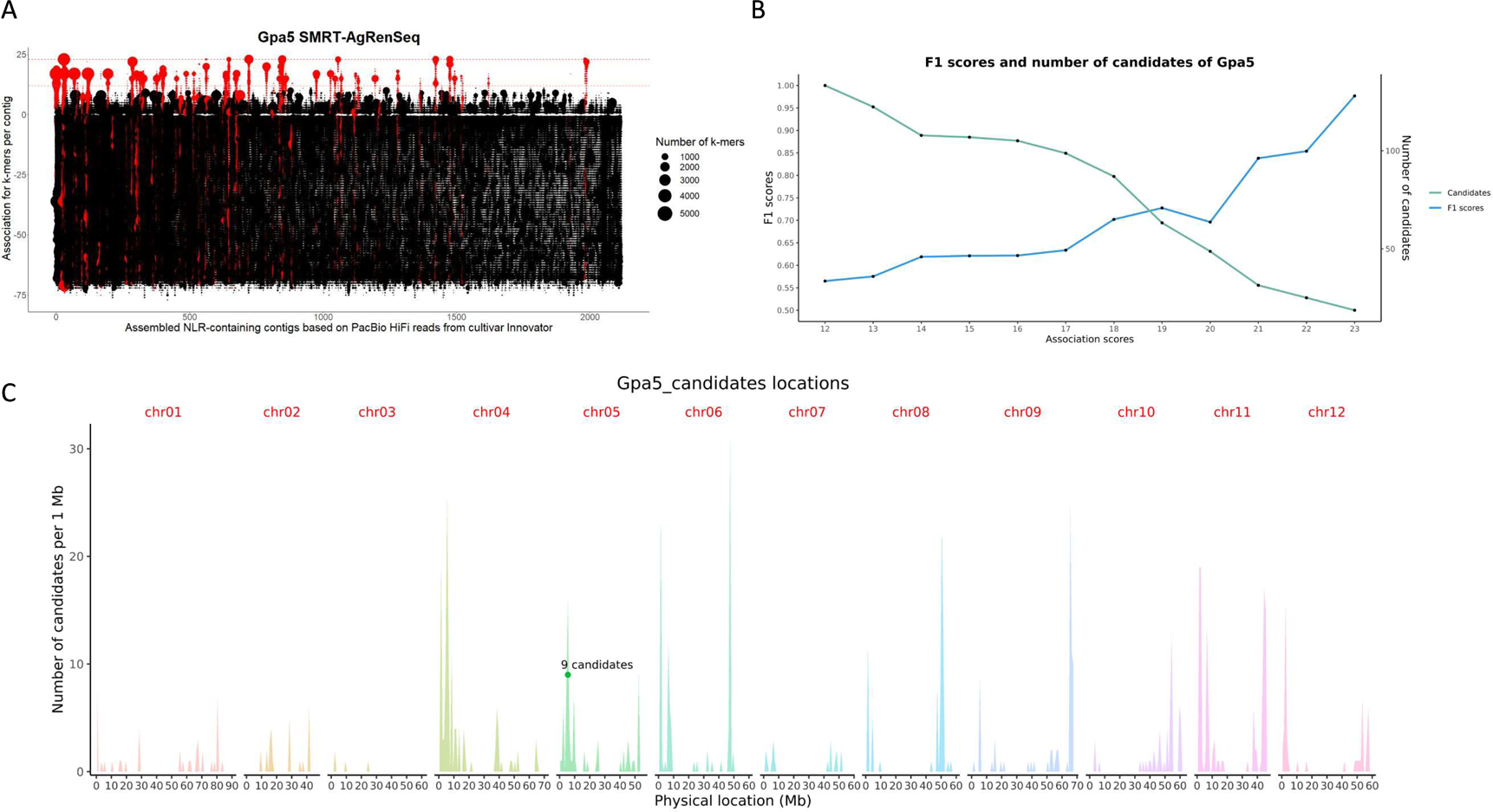
SMRT-AgRenSeq-d of *Gpa5*. a) Identification of NLR containing contigs in the reference cultivar Innovator with association to *Gpa5*. Columns on the X-axis represents contigs with signals of NLRs, with dots indicating mapped *k*-mers and their association score to *Gpa5* on the Y-axis. Dashed lines highlight highest association score (23, red) and chosen threshold (12, Orange). b) F1 score and number of candidate NLRs for each association score group. c) The predicted positions of *Gpa5* candidates with F1=1 are shown (dots) in relation to all NLRs from the reference DM v6.1 mapped over 12 chromosomes (peaks).

Following dRenSeq analysis, nine NLRs with an F1 value of 1 were identified and were considered strong candidates for the resistance (Supplementary Table 8). Consistent with Sattarzadeh *et al.* [11] and van Eck *et al.* [10] these nine candidates map to the *Rpi-R1* locus on potato Chromosome 5 (Figure 3c; Supplementary Table 8) and phylogenetically cluster with the *Rpi-R1* gene family (Figure 1).

Genetically, this study provides evidence that *Gpa5* is in repulsion to *Rpi-R1* as the *Rpi-R1* containing contig tig00000086 (Table 1) was not associated with the nematode resistance (Supplementary Table 8). This is further supported by the presence/absence of *Rpi-R1* in the association panel, where more than 60% (14 out of 23) of clones that contained *Gpa5* did not contain *Rpi-R1* (Supplementary Table 2). None of the candidates on Chromosome 9 passed the dRenSeq validation, albeit their F1 score was relatively high and ranged from 0.90 to 0.76 mainly due to their absence in some resistant varieties.

### *Gpa5* might require additional genes to provide full resistance

As detailed above, SMRT-AgRenSeq-d identified nine potential candidates that are strongly associated with *Gpa5*-based resistance. To further validate their role in providing resistance, we independently assessed the presence/absence of these candidate genes in three additional potato cultivars (Tiger, Novano and Karelia) that all yielded a resistance score of between 7 and 8 against *G. pallida* Pa2/3 (Supplementary Table 9). We further included five breeding clones that tested positive for the HC marker, but only displayed intermediate resistance alongside the cultivar Alicante that also provided an intermediate resistance (score of 6 against Pa3; 9 against Pa2). As a negative control, we included the cultivar Vales Everest which displayed intermediate to high resistance against *G. pallida* Pa2/3. However this cultivar utilises the *H3* resistance instead of *Gpa5* [31].

Except for Vales Everest, all candidates with an F1 score of 1 showed consistent presence in this set, including the lines with intermediate phenotypes. Interestingly, the NLRs from chromosome 9 were observed in members of the high and intermediate resistance scoring group, making them candidates for the added response. None of the *Gpa5* candidates were present in Vales Everest, which is consistent with *H3*, introduced from *S. tuberosum* ssp. *andigena* (CPC 2802), being a distinct resistance from the *S. vernei-*derived *Gpa5* [3,9].

Based on previous studies*, Gpa6* on chromosome 9 exhibits an additive effect with *Gpa5* in terms of resistance to *G. pallida* Pa2/3 [9]. The loss of *Gpa6* may account for the observed intermediate resistance in some clones in our panel whilst its absence in some resistant clones could have been masked by the presence of other functional genes such as *Grp1* or *H2*. Thus, the NLRs on chromosome 9 could represent potential *Gpa6* candidates (Supplementary Table 8) and warrant further studies in the future.

It is further conceivable that *Gpa5*, a potential *Rpi-R1* homolog (Figure 1), requires additional genes such as NRCs to function. Indeed, *Rpi-R1* requires NRC4 to elicit a functional response [28]. Thus, the intermediate resistance response observed in some potatoes, despite the presence of the candidate NLRs, could be a consequence of NRC variations that are less conducive to triggering a full response.

## Conclusion

In conclusion, we have further developed SMRT-AgRenSeq by including a dRenSeq analysis that significantly expedites the process of identifying candidates that are absolutely linked to the resistance phenotype. The F1 score that can be determined for all candidates in this combined approach is a highly informative measure of linkage and proved more informative than, for example, the association score alone. The resulting method, SMRT-AgRenSeq-d, successfully identified the benchmark genes *Rpi-R1, Rpi-R2-like, Rpi-R3a* and *Rpi-R3b* with the highest possible F1 score of 1 in the association panel. Importantly, SMRT-AgRenSeq-d correctly identified those genes even though their penetration in the panel varied between 8.5% for *Rpi-R2-like*, 29.3% for *Rpi-R1*, 37.6% for *Rpi-R3a*, and 51.3% for *Rpi-R3b*. When applied to *Gpa5,* which is currently the most effective and widespread resistance against *G. pallida* used in potato crop protection, SMRT-AgRenSeq-d identified strong candidate NLRs. All *Gpa5* candidates mapped to the previously established locus on potato chromosome 5 and were shown to be phylogenetically related to the late blight resistance gene *R1*. We propose that this refined method is suitable for studying NLR-based resistances in diverse crops and against diverse pathogens.

## Materials and Methods

### Source of plant and pathogen material

*G. pallida* cysts that were used in the phenotyping test were of the Pa2/3 population known as ‘Lindley’ [32]. The population was multiplied on the susceptible cultivar Desiree in glasshouses in 2019. All potato accessions were grown in the glasshouse. In some cases, lyophilised DNA was received from collaborators.

### Extraction of DNA

Bulk DNA was extracted from leaves of all potato accessions using the Qiagen DNEasy Plant Mini Kit (Qiagen, Hilden, Germany) following the manufacturer’s instructions. DNA for HiFi sequencing was extracted using the Wizard® HMW DNA Extraction Kit (Promega, Madison, WI, USA), following the manufacturer’s instructions.

### RenSeq

The RenSeq bait library was as described in Armstrong *et al.* [20]. Enrichment sequencing was performed by Arbor Biosciences (Ann Arbor, MI, USA) using the Novaseq 6000 platform for Illumina reads and the PacBio Sequel II platform for HiFi reads.

### Assembly

Raw Illumina reads were mapped to the DM v6.1 genome assembly [33] with BWA-mem v0.7.17-r1188 [34] using default parameters. Picard CollectInsertSizeMetrics v2.25.1 [35] was used to calculate the mean and standard deviation of the read insert size. This was used to inform the MASURCA [36] v3.4.2 assembly with PE= pe 342 97 and CLOSE_GAPS=1 as non-default parameters. Assembly of HiFi reads was conducted using HiCanu with the SMRT-RenSeq Assembly workflow within HISS v2.0.0 [37,38].

### Identification of known resistance genes

The CDS sequences of *R1, R2-like, R3a* and *R3b* [39–42] were searched for in both sets of assembled contigs using BLASTN v2.11.0 [43,44] with an e-value of 1e-5.

### Phylogenetics

NLR sequences from the SMRT-RenSeq assemblies were predicted using NLR-Annotator [25] and Nucleotide Binding (NB) domains were identified with Interproscan version 5.54-87.0 [45,46] with goterms, iprlookup and pathways options enabled. The output XML file was parsed with a custom python script [47] to extract the amino acid sequences of the NB domains (IPR002182). Reference NRC amino acid sequences were retrieved from Adachi *et al.* [48] and submitted to Interproscan and the output tsv file was parsed with a different custom python script [47]. The BED files produced by these scripts were used to extract FASTA sequences with the bedtools getfasta function (version 2.30.0) [49]. The known NLR and NRC gene NB domain sequences were aligned with Clustal Omega version 1.2.4 [50] with 10 iterations. Following this, the NB domains of the automatically predicted NLRs were added to the alignment with Clustal Omega using the previous alignment as a profile, again with 10 iterations.

Tree construction was performed in R v4.1.3 [51] using the libraries ape v5.6-2 [52] and phangorn v2.9.0 [53]. Sites and sequences with 85% missing data, or more, were removed from the alignment. Following this, duplicated sequences were removed from the alignment. A phylogeny based on maximum likelihood was inferred from the final alignment. The Bayesian information criteria (BIC) obtained from the “modelTest” function of phangorn suggested a JTT+G model. The tree was bootstrapped with 1,000 replicates and its topology was optimised using nearest neighbour interchanges (NNI). The phylogeny produced was then rooted to the outgroup sequences from *Homo sapiens* and *Caenorhabditis elegans*. Clades were assigned with PhyloPart 2.1 [54] with a percentile threshold of 0.05. The figure was created in IToL [55], details on how to replicate the figure and text files for modifying the view are described in a markdown file [47].

### Diversity panel formation and additional phenotyping

The resistance or susceptibility of clones within the association panel was determined by retrieving results from potato cultivar databases and/or through replicated PCN tests. Only resistant clones that also tested positive for the HC marker [11] were considered to ensure that the source of the resistance is consistent (Supplementary Table 3; Supplementary Table 9). The phenotyping experiment was based on the method published by [56]. Briefly, tubers of breeding clones and six controls (Desiree, Maris Piper, Vales Everest, Lady Balfour, Innovator and King Russet) were planted in canisters (one tuber per canister) with four replicates. All canisters were placed into four trays, each containing one replicate of all breeding clones and controls in a randomized design. Canisters were inoculated with approximately 10–15 cysts. After inoculation, canisters were stored in the dark at 18°C for seven weeks. After this time, the number of females observed on the root balls were counted, and the phenotypic scores were recorded. In these accessions and breeding clones, 117 lines whose scores were between 1 and 3 (susceptible accessions) and 7–9 (resistant accessions) were chosen for the diversity panel for *Gpa5*.

To determine the presence/absence of the late blight resistance genes *Rpi-R1, Rpi-R2-like, Rpi-R3a* and *Rpi-R3b* that are present in Innovator, dRenSeq was performed as described previously by Armstrong *et al.* [20]. Based on the known diversity within the *Rpi-R2* family [20], we set a minimum sequence coverage threshold of 97% for a gene to be classified as present.

### SMRT-AgRenSeq-d

SMRT-AgRenSeq-d was performed using the AgRenSeq and dRenSeq workflows in HISS v2.0.0 [37]. AgRenSeq followed the workflow initially described for *A*. *tauschii* [22]. Briefly, the Illumina reads of the previously described panel of samples were assessed for the presence and absence of *k*-mers with a *k* value of 51 and arranged into a presence/absence matrix. This matrix was then used alongside phenotypic information to perform the association and the identities of contigs containing *k*-mers highly associated with the phenotype were reported.

An initial threshold of 50% of the highest association score was used for selecting candidate contigs and the NLR coordinates were determined with NLR-Annotator [25] (*Rpi-R1*: 17; *Rpi-R2-like*: 5; *Rpi-R3a*: 22; *Rpi-R3b*: 30; *Gpa5*: 12). These selected contigs were used for dRenSeq. This followed the pipeline previously described [20,57] whilst specifically reporting on NLR regions linked to bait coverage. Briefly, the Illumina reads of the panel were mapped to the CDS regions of candidate genes. This mapping was then filtered to contain only reads with zero mismatches within regions also represented by baits with at least 90% sequence identity. The resulting coverage of the candidates was reported.

### Refining of candidates

Candidates identified by SMRT-AgRenSeq-d were further refined through the generation of precision and recall values and calculation of the F1 score of the candidates [30]. The following formula was applied:

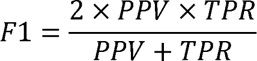

PPV is the Positive Predictive Value or precision, calculated by the following formula:

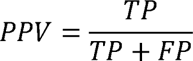

TPR is the True Positive Rate or sensitivity/recall, calculated by the following formula:

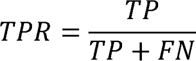

Here, TP represents true positive cases i.e., varieties/clones that are phenotypically resistant and have complete coverage of the candidate in dRenSeq (within the parameters specified above). FP (false positive) refers to varieties that are phenotypically susceptible but still had complete coverage in dRenSeq. Conversely, TN (true negative) refers to varieties that are phenotypically susceptible and had only partial coverage in dRenSeq, whereas FN (false negative) refers to phenotypically resistant varieties with partial coverage in dRenSeq. The individual F1 scores were calculated for every candidate.

## Acknowledgements

The authors thank John Jones for providing critical feedback on the manuscript. Further we acknowledge the Research/Scientific Computing teams at The James Hutton Institute and NIAB for providing computational resources and technical support for the “UK’s Crop Diversity Bioinformatics HPC” (BBSRC grant BB/S019669/1), use of which has contributed to the results reported within this paper.

This work was supported by the Rural & Environment Science & Analytical Services (RESAS) Division of the Scottish Government through project JHI-B1-1, the Biotechnology and Biological Sciences Research Council (BBSRC) through award BB/S015663/1 and the Royal Society through award NAF\R1\201061. YW was supported through the CSC scholarship program, China. LB was supported through the East of Scotland Bioscience Doctoral Training Partnership (EASTBIO DTP), funded by the BBSRC award BB/T00875X/1. AK was supported through a Research Leaders 2025 fellowship funded by European Union’s Horizon 2020 research and innovation programme under Marie Sklodowska-Curie grant agreement no. 754380.

## Contributions

YW, LB, TMA, MA, AK conducted the computational analyses, developed the association panel and the F1 score. JL, XW, BH provided material for this study and contributed towards the writing of the manuscript and the independent validation. VY, DT and YW conducted the PCN score and HC marker analysis. TMA and KN conducted the phylogenetic analysis. SO provided genomic data and recombination analysis for this study. MB and IH conceived this work and secured funding. All authors have read and endorsed the manuscript.

## Data availability statement

BioProjects:

ERP141787 – Illumina Reads / contigs

ERP141789 – HiFi Reads / contigs

## Supplementary Tables

Supplementary Table 1: Read and assembly statistics for RenSeq and SMRT-RenSeq derived sequence presentation of the NLRome from the cultivar Innovator.

Supplementary Table 2: Description of the association panel. Highlighted are the cultivars and breeding clones used in this study and their coverage of the benchmark genes *Rpi-R1, Rpi-R2-like, Rpi-R3a* and *Rpi-R3b* as established through dRenSeq. Coverage exceeding 97% of the full-length gene equates to the presence of the gene. Presence/absence is transposed into a +1/-1 score used for the association study. For *Gpa5*, only cultivars that yielded a disease resistance score between 7 and 9 or 1–3, were used to represent resistant/susceptible, respectively. Further, all resistant plants had to also yield a positive PCR test for the HC marker to ensure the source of the resistance is consistent. ENA read accession numbers are highlighted for all plants in the panel.

Supplementary Table 3: Immediate descendent potato cultivars and breeding clones from the cultivar Innovator (URL: http://www.plantbreeding.wur.nl/PotatoPedigree/). Shown are the parents of the cross, the year, breeder and breeder code.

Supplementary Table 4: dRenSeq analysis of *Rpi-R3a.* Depicted is the dRenSeq-based coverage of NLR candidates with an association score of larger than 22 across 117 potato clones representing the association panel. Based on the presence/absence of candidate NLRs in potato clones associated with the *Rpi-R3a* and those that do not contain *Rpi-R3a*, an F1 score was calculated. Only candidates with an F1 score of 1 display absolute linkage to the phenotype.

Supplementary Table 5: dRenSeq analysis of *Rpi-R3b.* Depicted is the dRenSeq-based coverage of NLR candidates with an association score of larger than 30 and an F1 score of larger than 0.7 across 117 potato clones representing the association panel. The F1 score was calculated based on the presence/absence of candidate NLRs in potato clones associated with the *Rpi-R3b* and those that do not contain *Rpi-R3b*.

Supplementary Table 6: dRenSeq analysis of *Rpi-R2-like*. Depicted is the dRenSeq-based coverage of NLR candidates with an association score of larger than 5 and an F1 score of larger than 0.7 across 117 potato clones representing the association panel. The F1 score was calculated based on the presence/absence of candidate NLRs in potato clones associated with the *Rpi-R2-like* and those that do not contain *Rpi-R2-like*.

Supplementary Table 7: dRenSeq analysis of *Rpi-R1*. Depicted is the dRenSeq-based coverage of NLR candidates with an association score of larger than 17 and an F1 score of larger than 0.7 across 116 potato clones representing the association panel. The F1 score was calculated based on the presence/absence of candidate NLRs in potato clones associated with the *Rpi-R1* and those that do not contain *Rpi-R1*.

Supplementary Table 8: dRenSeq analysis of *Gpa5.* Depicted is the dRenSeq-based coverage of NLR candidates with an association score of larger than 12 and an F1 score of larger than 0.7 across 117 potato clones representing the association panel. The F1 score was calculated based on the presence/absence of candidate NLRs in potato clones associated with the *Gpa5* and those that do not contain *Gpa5*.

Supplementary Table 9: dRenSeq analysis of *Gpa5* candidate NLRs in additional potato clones with high and intermediate resistance against *Globodera pallida* Pa2/3. The cultivar Vales Everest is used as a negative control since the resistance is known to be independent of *Gpa5* and utilises instead *H3*.

## Supplementary Figures

Supplementary Figure 1: Phylogenetic tree displaying NB domains of Innovator NLRs following assembly of SMRT-RenSeq reads. All clades are expanded. Highlighted are the identified benchmark genes and candidates from each association with F1=1. The NB containing proteins APAF1 from *Homo sapiens* and CED4 from *Caenorhabditis elegans* act as outgroups. Bootstrap values are displayed as branch labels. The tree scale indicates substitutions/site.

Supplementary Figure 2: SMRT-AgRenSeq-d analysis for *Rpi-R3b*. a) Columns on the X-axis represents NLR containing contigs, with dots indicating mapped *k*-mers and their association score to *Rpi-R3b* on the Y-axis. Dashed lines highlight highest association score (60, red) and chosen threshold (30, Orange). b) F1 score and the number of candidate NLRs for each association score group. c) The predicted position of *Rpi-R3b* candidates with F1=1 are shown (dots) in relation to all NLRs from the reference DM v6.1 mapped over 12 chromosomes (peaks).

Supplementary Figure 3: SMRT-AgRenSeq-d analysis for *Rpi-R2-like*. a) Columns on the X-axis represents NLR containing contigs, with dots indicating mapped *k*-mers and their association score to *Rpi-R2-like* on the Y-axis. Dashed lines highlight highest association score (10, red) and chosen threshold (5, Orange). b) F1 score and the number of candidate NLRs for each association score group. c) The predicted position of *Rpi-R2-like* candidates with F1=1 are shown (dots) in relation to all NLRs from the reference DM v6.1 mapped over 12 chromosomes (peaks).

Supplementary Figure 4: SMRT-AgRenSeq-d analysis for *Rpi-R1*. a) Columns on the X-axis represents NLR containing contigs, with dots indicating mapped *k*-mers and their association score to *Rpi-R1* on the Y-axis. Dashed lines highlight highest association score (34, red) and chosen threshold (17, Orange). b) F1 score and the number of candidate NLRs for each association score group. c) The predicted position of *Rpi-R1* candidates with F1=1 are shown (dots) in relation to all NLRs from the reference DM v6.1 mapped over 12 chromosomes (peaks).

## References

1. Birch PRJ, Bryan G, Fenton B, et al. Crops that feed the world 8: Potato: are the trends of increased global production sustainable? Food Sec 2012;4:477–508.

2. Tiwari JK, Buckseth T, Zinta R et al. Germplasm, Breeding, and Genomics in Potato Improvement of Biotic and Abiotic Stresses Tolerance. Frontiers in Plant Science 2022;13.

3. Gartner U, Hein I, Brown LH et al. Resisting Potato Cyst Nematodes With Resistance. Frontiers in Plant Science 2021;12.

4. Jones JT, Haegeman A, Danchin EGJ et al. Top 10 plant-parasitic nematodes in molecular plant pathology. Molecular Plant Pathology 2013;14:946–61.

5. Turner SJ. Population decline of potato cyst nematodes (Globodera rostochiensis, G. pallida) in field soils in Northern Ireland. Annals of Applied Biology 1996;129:315– 22.

6. Price JA, Coyne D, Blok VC et al. Potato cyst nematodes Globodera rostochiensis and G. pallida. Molecular Plant Pathology 2021;22:495–507.

7. Paal J, Henselewski H, Muth J et al. Molecular cloning of the potato Gro1-4 gene conferring resistance to pathotype Ro1 of the root cyst nematode Globodera rostochiensis, based on a candidate gene approach. The Plant Journal 2004;38:285–97.

8. Van Der Vossen EAG, Van Der Voort JNAMR, Kanyuka K et al. Homologues of a single resistance-gene cluster in potato confer resistance to distinct pathogens: a virus and a nematode. The Plant Journal 2000;23:567–76.

9. Rouppe van der Voort J, van der Vossen E, Bakker E et al. Two additive QTLs conferring broad-spectrum resistance in potato to Globodera pallida are localized on resistance gene clusters: Theor Appl Genet 2000;101:1122–30.

10. van Eck HJ, Vos PG, Valkonen JPT et al. Graphical genotyping as a method to map Ny(o,n)sto and Gpa5 using a reference panel of tetraploid potato cultivars. Theor Appl Genet 2017;130:515–28.

11. Sattarzadeh A, Achenbach U, Lübeck J et al. Single nucleotide polymorphism (SNP) genotyping as basis for developing a PCR-based marker highly diagnostic for potato varieties with high resistance to Globodera pallida pathotype Pa2/3. Mol Breeding 2006;18:301–12.

12. Finkers-Tomczak A, Danan S, van Dijk T et al. A high-resolution map of the Grp1 locus on chromosome V of potato harbouring broad-spectrum resistance to the cyst nematode species Globodera pallida and Globodera rostochiensis. Theor Appl Genet 2009;119:165–73.

13. Bryan G, McLean K, Bradshaw J et al. Mapping QTLs for resistance to the cyst nematode Globodera pallida derived from the wild potato species Solanum vernei. Theor Appl Genet 2002;105:68–77.

14. Dalton E, Griffin D, Gallagher TF et al. The effect of pyramiding two potato cyst nematode resistance loci to Globodera pallida Pa2/3 in potato. Mol Breeding 2013;31:921–30.

15. Jupe F, Witek K, Verweij W et al. Resistance gene enrichment sequencing (RenSeq) enables reannotation of the NB-LRR gene family from sequenced plant genomes and rapid mapping of resistance loci in segregating populations. The Plant Journal 2013;76:530–44.

16. Jupe F, Pritchard L, Etherington GJ et al. Identification and localisation of the NB-LRR gene family within the potato genome. BMC Genomics 2012;13:75.

17. Xu X, Pan S, Cheng S et al. Genome sequence and analysis of the tuber crop potato. Nature 2011;475:189–95.

18. Van de Weyer A-L, Monteiro F, Furzer OJ et al. A Species-Wide Inventory of NLR Genes and Alleles in Arabidopsis thaliana. Cell 2019;178:1260–1272.

19. Witek K, Jupe F, Witek AI et al. Accelerated cloning of a potato late blight– resistance gene using RenSeq and SMRT sequencing. Nature Biotechnology 2016;34:656–60.

20. Armstrong MR, Vossen J, Lim TY et al. Tracking disease resistance deployment in potato breeding by enrichment sequencing. Plant Biotechnology Journal 2019;17:540–9.

21. Chen X, Lewandowska D, Armstrong MR et al. Identification and rapid mapping of a gene conferring broad-spectrum late blight resistance in the diploid potato species Solanum verrucosum through DNA capture technologies. Theoretical and Applied Genetics 2018;131:1287–97.

22. Arora S, Steuernagel B, Gaurav K et al. Resistance gene cloning from a wild crop relative by sequence capture and association genetics. Nat Biotechnol 2019;37:139–43.

23. Vendelbo NM, Mahmood K, Steuernagel B et al. Discovery of Resistance Genes in Rye by Targeted Long-Read Sequencing and Association Genetics. Cells 2022;11:1273.

24. Corrion A, Day B. Pathogen Resistance Signalling in Plants. ELS. John Wiley & Sons, Ltd, 2015, 1–14.

25. Steuernagel B, Witek K, Krattinger SG et al. The NLR-Annotator Tool Enables Annotation of the Intracellular Immune Receptor Repertoire. Plant Physiol 2020;183:468–82.

26. Baumgarten A, Cannon S, Spangler R et al. Genome-Level Evolution of Resistance Genes in Arabidopsis thaliana. Genetics 2003;165:309–19.

27. Huang Z, Qiao F, Yang B et al. Genome-wide identification of the NLR gene family in Haynaldia villosa by SMRT-RenSeq. BMC Genomics 2022;23:118.

28. Wu C-H, Abd-El-Haliem A, Bozkurt TO et al. NLR network mediates immunity to diverse plant pathogens. Proc Natl Acad Sci U S A 2017;114:8113–8.

29. van Berloo R, Hutten RCB, van Eck HJ, et al. An Online Potato Pedigree Database Resource. Potato Res 2007;50:45–57.

30. Chinchor N, Diego S. MUC-4 EVALUATION METRICS. 1992, 22–9.

31. Adillah Tan MY, Park T-H, Alles R et al. GpaXItarloriginating from Solanum tarijense is a major resistance locus to Globodera pallida and is localised on chromosome 11 of potato. Theor Appl Genet 2009;119:1477–87.

32. Kaczmarek AM, Back M, Blok VC. Population dynamics of the potato cyst nematode, Globodera pallida, in relation to temperature, potato cultivar and nematicide application. Plant Pathology 2019;68:962–76.

33. Pham GM, Hamilton JP, Wood JC et al. Construction of a chromosome-scale long-read reference genome assembly for potato. GigaScience 2020;9, DOI: 10.1093/gigascience/giaa100.

34. Li H. Aligning sequence reads, clone sequences and assembly contigs with BWA-MEM. 2013, DOI: 10.48550/arXiv.1303.3997.

35. Broad Institute,. “Picard Toolkit.” GitHub repository https://broadinstitute.github.io/picard/, Broad Institute 2019.

36. Zimin AV, Marçais G, Puiu D et al. The MaSuRCA genome assembler. Bioinformatics 2013;29:2669–77.

37. Adams T, Smith M, Bayer M et al. HISS: Snakemake-based workflows for performing SMRT-RenSeq assembly, AgRenSeq and dRenSeq for the discovery of novel plant disease resistance genes. 2022:2022.11.01.514708.

38. Nurk S, Walenz BP, Rhie A et al. HiCanu: accurate assembly of segmental duplications, satellites, and allelic variants from high-fidelity long reads. Genome Res 2020;30:1291–305.

39. Ballvora A, Ercolano MR, Weiss J et al. The R1 gene for potato resistance to late blight (Phytophthora infestans) belongs to the leucine zipper/NBS/LRR class of plant resistance genes. Plant J 2002;30:361–71.

40. Huang S, Van Der Vossen EAG, Kuang H et al. Comparative genomics enabled the isolation of the R3a late blight resistance gene in potato. The Plant Journal 2005;42:251–61.

41. Li G, Huang S, Guo X et al. Cloning and Characterization of R3blJ; Members of the R3 Superfamily of Late Blight Resistance Genes Show Sequence and Functional Divergence. MPMI 2011;24:1132–42.

42. Lokossou AA, Park T, van Arkel G, et al. Exploiting Knowledge of R/Avr Genes to Rapidly Clone a New LZ-NBS-LRR Family of Late Blight Resistance Genes from Potato Linkage Group IV. MPMI 2009;22:630–41.

43. Altschul SF, Gish W, Miller W et al. Basic local alignment search tool. J Mol Biol 1990;215:403–10.

44. Zhang Z, Schwartz S, Wagner L et al. A greedy algorithm for aligning DNA sequences. J Comput Biol 2000;7:203–14.

45. Blum M, Chang H-Y, Chuguransky S et al. The InterPro protein families and domains database: 20 years on. Nucleic Acids Research 2021;49:D344–54.

46. Jones P, Binns D, Chang H-Y et al. InterProScan 5: genome-scale protein function classification. Bioinformatics 2014;30:1236–40.

47. Adams T, Wang Y, Wangand K. TMAdams/gpa5_smrt_agrenseq_paper: v1.0.0. 2022.

48. Adachi H, Sakai T, Harant A, et al. An atypical NLR protein modulates the NRC immune receptor network. 2021:2021.11.15.468391.

49. Quinlan AR, Hall IM. BEDTools: a flexible suite of utilities for comparing genomic features. Bioinformatics 2010;26:841–2.

50. Sievers F, Wilm A, Dineen D et al. Fast, scalable generation of high-quality protein multiple sequence alignments using Clustal Omega. Molecular Systems Biology 2011;7:539.

51. R Core Team. R: A language and environment for statistical computing. R: A language and environment for statistical computing. R Foundation for Statistical Computing, Vienna, Austria. URL https://www.R-project.org/. 2022.

52. Paradis E, Schliep K. ape 5.0: an environment for modern phylogenetics and evolutionary analyses in R. Bioinformatics 2019;35:526–8.

53. Schliep KP. phangorn: phylogenetic analysis in R. Bioinformatics 2011;27:592–3.

54. Prosperi MCF, Ciccozzi M, Fanti I et al. A novel methodology for large-scale phylogeny partition. Nat Commun 2011;2:321.

55. Letunic I, Bork P. Interactive Tree Of Life (iTOL) v5: an online tool for phylogenetic tree display and annotation. Nucleic Acids Research 2021;49:W293–6.

56. Phillips MS, Forrest JMS, Wilson LA. Screening for resistance to potato cyst nematode using closed containers. Annals of Applied Biology 1980;96:317–22.

57. Van Weymers PSM, Baker K, Chen X et al. Utilizing “Omic” Technologies to Identify and Prioritize Novel Sources of Resistance to the Oomycete Pathogen Phytophthora infestans in Potato Germplasm Collections. Front Plant Sci 2016;7, DOI: 10.3389/fpls.2016.00672.

