## Supplementary figures and images for "SMRT-AgRenSeq-d in potato (Solanum tuberosum) identifies candidates for the nematode resistance Gpa5"

### Supplementary Figure 1

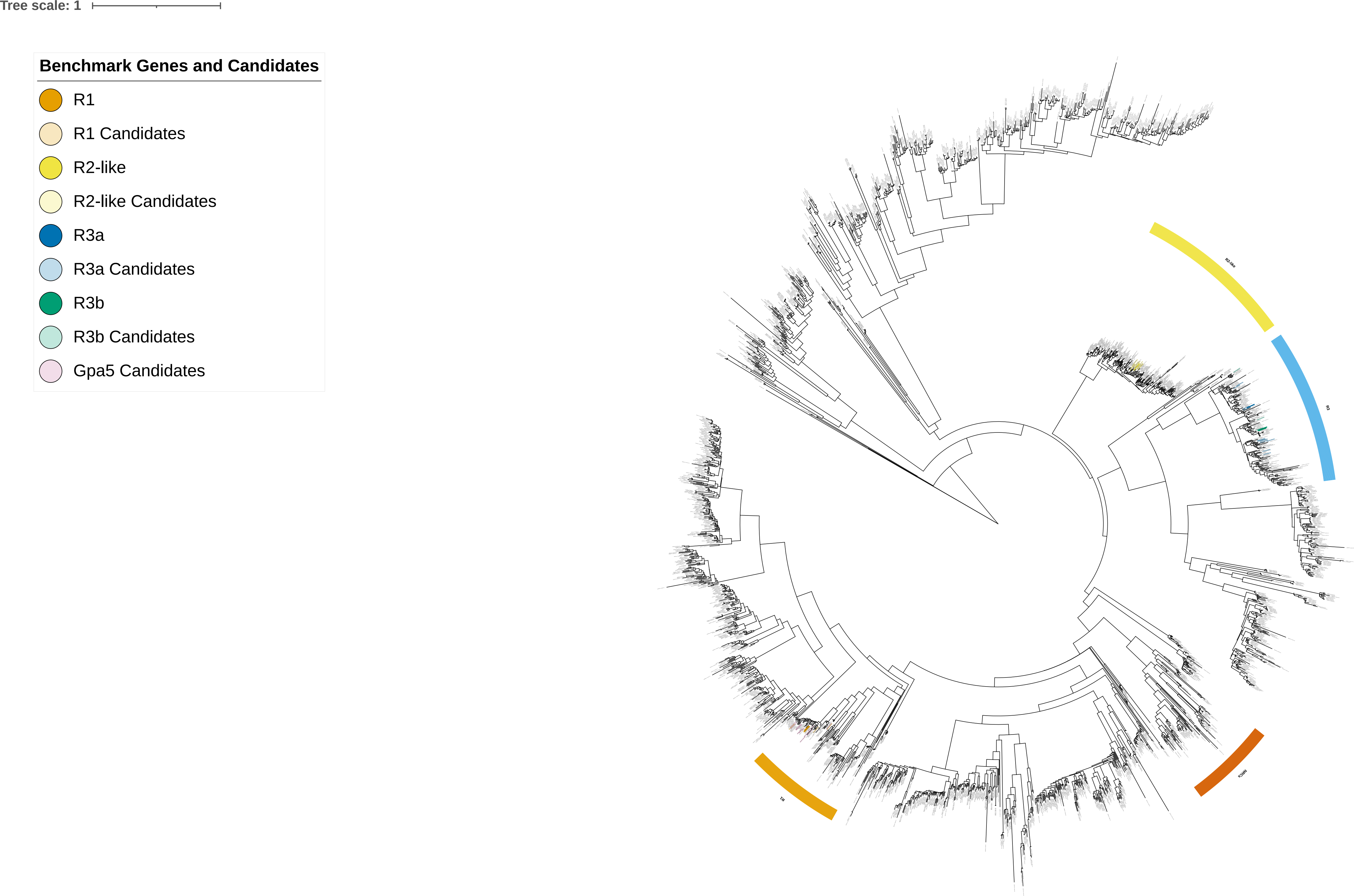

### Supplementary Figure 2

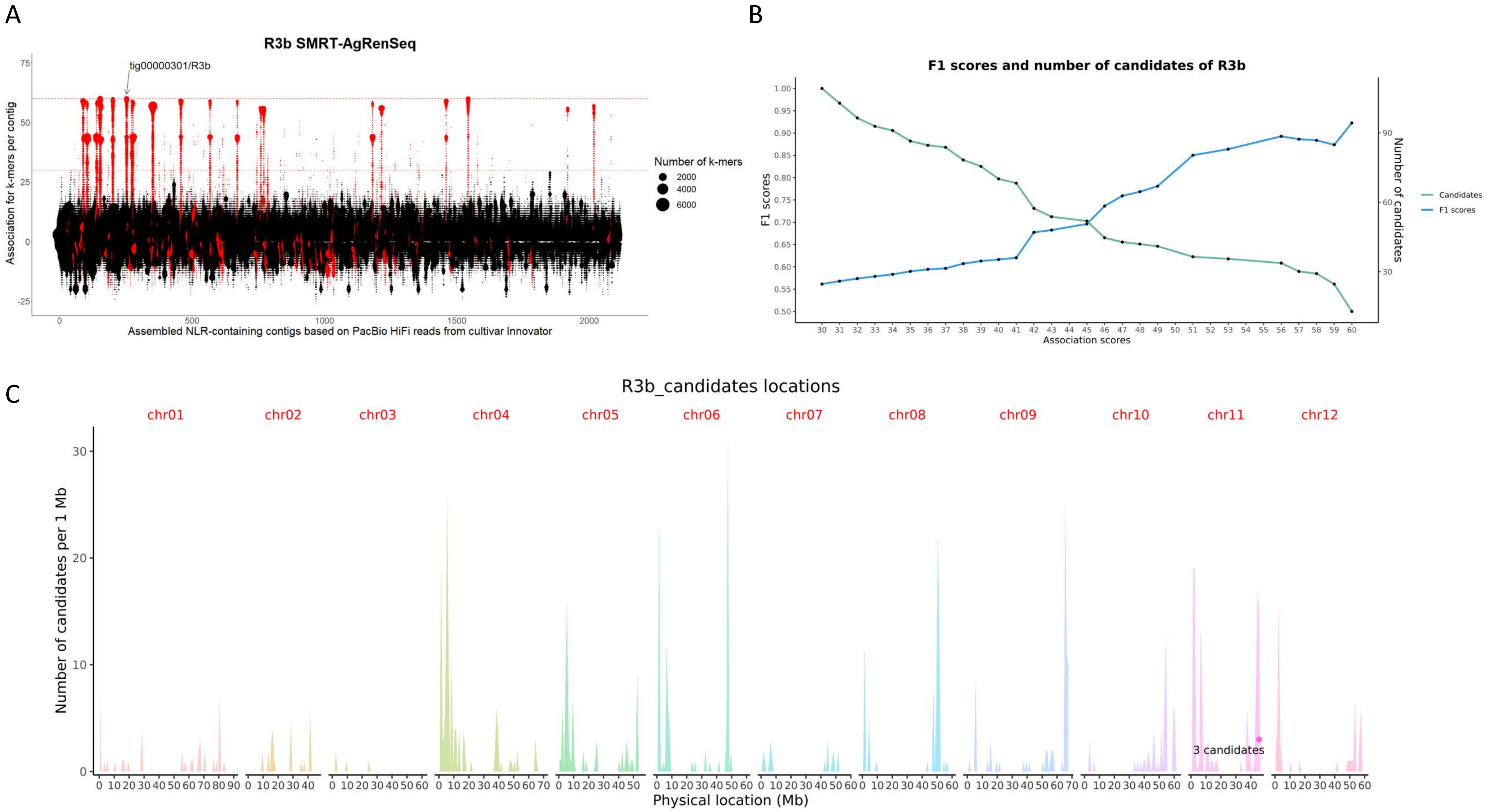

### Supplementary Figure 3

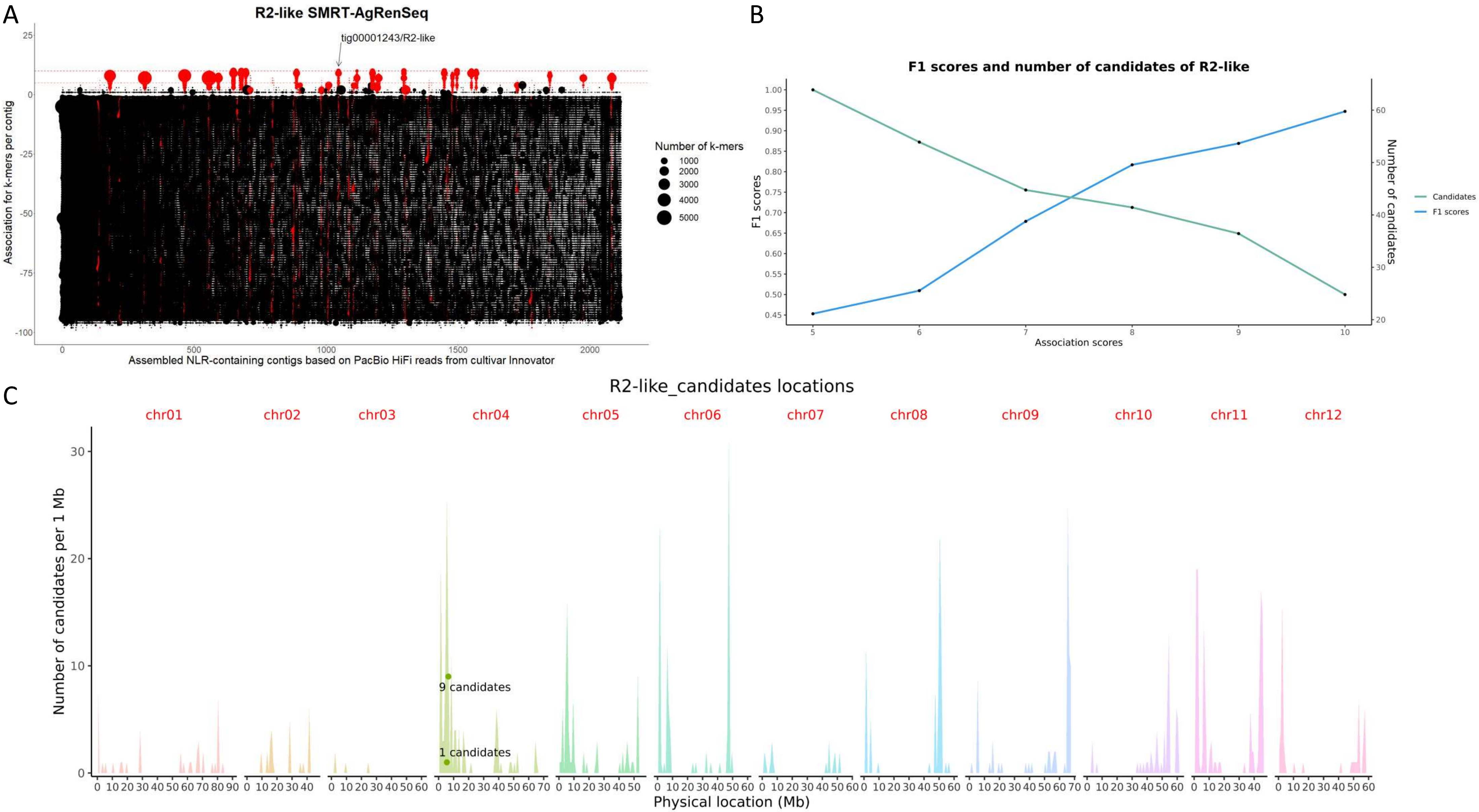

### Supplementary Figure 4

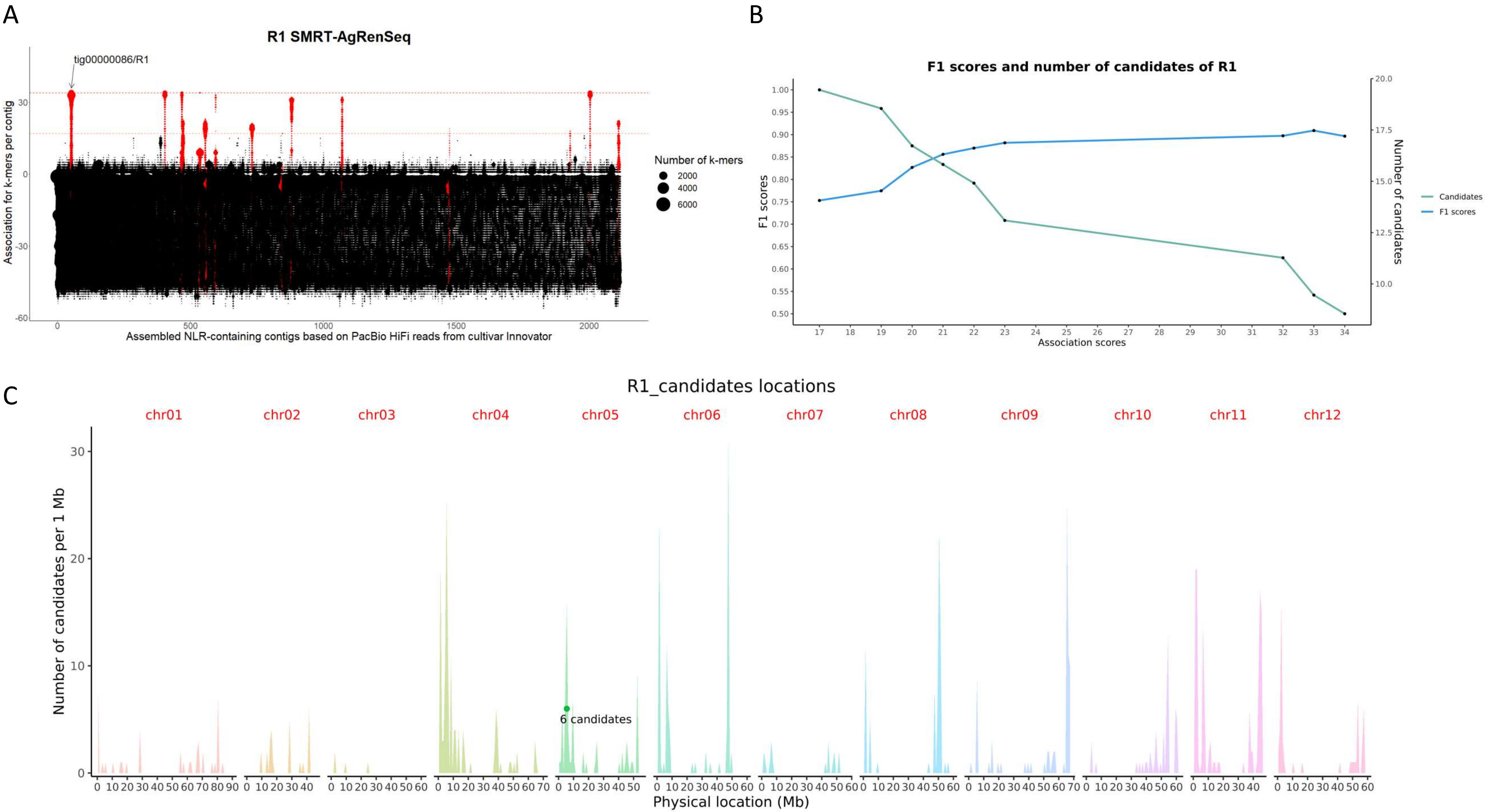
